# Neural tracking of auditory statistical regularities is reduced in adults with dyslexia

**DOI:** 10.1101/2024.08.26.609678

**Authors:** Hanna Ringer, Daniela Sammler, Tatsuya Daikoku

**Author notes:** Corresponding author: Dr. Hanna Ringer, The University of Tokyo, 7-3-1 Hongo, Bunkyo-ku, Tokyo 113-8656, Japan.

## Abstract

Listeners implicitly use statistical regularities to segment continuous sound input into meaningful units, e.g., transitional probabilities between syllables to segment a speech stream into separate words. Implicit learning of such statistical regularities in a novel stimulus stream is reflected in a synchronisation of neural responses to the sequential stimulus structure. The present study aimed to test the hypothesis that neural tracking of the statistical stimulus structure is reduced in individuals with dyslexia who have weaker reading and spelling skills, and possibly also weaker statistical learning abilities in general, compared to healthy controls. To this end, adults with and without dyslexia were presented with continuous streams of (non-speech) tones, which were arranged into triplets, such that transitional probabilities between single tones were high within triplets and low between triplets. We found that neural tracking of the triplet structure, i.e., phase coherence of EEG response at the triplet rate relative to the tone rate, was reduced in adults with dyslexia compared to the control group. Moreover, enhanced neural tracking of the statistical structure was associated with better spelling skills. These results suggest that individuals with dyslexia have a rather broad deficit in processing structure in sound instead of a merely phonological deficit.

## Introduction

One of the key challenges of speech perception is to segment the continuous speech signal into meaningful units, such as words, syllables, or phonemes. Acquiring implicit knowledge that helps to accomplish this task plays a particularly crucial role during language learning, both during early childhood and when learning a second language later in life. Difficulties in acquiring and applying such knowledge to structure speech input, or acoustic input more generally, may be detrimental for the efficiency of speech perception, and therefore communication and well-being. In sequentially structured sound materials such as speech or music, short- and long-range statistical relationships that govern the order in which single elements (e.g., syllables or tones) are arranged within a longer sequence (e.g., words or melodic phrases) are a critical cue for the segmentation of acoustic input. In particular, segmentation occurs based on transitional probabilities between single elements, i.e., conditional probabilities for particular items to follow specific preceding items (Daikoku 2018). Sensitivity to the transitional probabilities between elements within a sound sequence is acquired through a powerful learning mechanism referred to as statistical learning (Saffran 2003). When presented with a constant stream of syllables that consisted of four trisyllabic artificial “words” repeated in a random order, 8-months-old infants rapidly learnt to segment the continuous syllable stream into distinct words based on transitional probabilities, which were high within words and low between words (Saffran et al. 1996). Automatic segmentation of stimulus streams based on transitional probabilities was subsequently also found for non-linguistic stimulus material in different sensory modalities, including sequences of pure tones (Saffran et al. 1999), visually presented shapes (Turk-Browne et al. 2005; Henin et al. 2021), and tactile vibration pulses at different stimulation sites (Conway and Christiansen 2005). Similar patterns of results were observed across different age groups from neonates (Teinonen et al. 2009; Bulf et al. 2011) to children and adults (Saffran et al. 1997), suggesting that statistical learning constitutes an innate capacity of the human brain. Moreover, it occurred largely implicitly, i.e., through passive exposure without explicit awareness that learning took place and without active access to what was learnt (Perruchet and Pacton 2006; Conway 2020), though some evidence suggests that learning of artificial words can be enhanced through explicit learning tasks, e.g., by providing linguistic information about the words such as written word forms (Chen et al. 2020).

At the neural level, perceptual segmentation of continuous acoustic streams based on their inherent (statistical) structure is achieved via neural tracking of the structure of the ongoing stimulation, i.e., the synchronisation of oscillatory neural activity with the stimulation, sometimes referred to as neural entrainment (for a recent review on neural tracking as a measure of statistical learning in syllable streams, see Sjuls et al. 2023). Functionally, this mechanism was interpreted as the alignment of peaks in rhythmically fluctuating attention and perceptual sensitivity to the most relevant time windows within a continuous stimulus (Jones 1976; Large and Jones 1999; Henry and Herrmann 2014; Riess Jones 2019). The frequency-tagging approach builds upon the tendency of the brain to automatically track periodically recurring events in continuous auditory stimulation and allows to experimentally test which perceptual units the brain responds to. Previous research using electroencephalography (EEG) combined with an adapted version of the statistical learning paradigm introduced above (Saffran et al. 1996) demonstrated that listeners’ brain activity synchronised with the rate of the trisyllabic words, which were separated by subliminal 25-ms pauses in structured syllable streams (Buiatti et al. 2009). More specifically, a peak in spectral power emerged at the word rate, whereas the peak at the syllable rate decreased while participants listened to the structured syllable streams (compared to random streams without a word structure; Buinatti et al. 2009). Strikingly, subsequent studies showed the same pattern of results even when trisyllabic words were not separated by pauses, i.e., when there were no acoustic cues in the syllable stream and segmentation must occur solely based on transitional probabilities between syllables (Batterink and Paller 2017; Batterink and Paller 2019; Choi et al. 2020). The so-called Word Learning Index (WLI), reflecting the strength of neural tracking (in terms of inter-trial coherence (ITC) of neural activity) at the word rate relative to the syllable rate, increased most strongly during the first few hundred word presentations. This suggests that sensitivity to the word structure in the syllable stream increased rapidly through passive exposure (Batterink and Choi 2021). Statistical learning of the word structure, as indexed by an increase of the WLI, was largely independent of listeners’ attentional focus (Batterink and Paller 2019) and occurred in both adults (Batterink and Paller 2017) and infants (Choi et al. 2020) alike. Concurrent tracking of syllables and words during statistical word learning was observed both in the presence and absence of additional prosodic cues, such as melodic contours (François et al. 2017) or stress patterns (Elmer et al. 2021) that corresponded to word boundaries. Other studies demonstrated robust neural tracking not only for syllables and words, but also for linguistic structures at longer time scales such as phrases or sentences (Ding et al. 2016), which was assumed to support parsing of the hierarchical syntactic structure of an utterance (Kazanina and Tavano 2023). Furthermore, a similar pattern of changes in neural tracking was observed for structured (compared to random) sequences of tones, suggesting that statistical learning shares similar neural signatures across different types of auditory stimulus material (Farthouat et al. 2017; Moser et al. 2021). Together, these results suggest that neural tracking does not merely reflect sensory processing of acoustic features, but automatic hierarchical integration of single elements within a sequence into meaningful perceptual units.

A growing body of research has suggested that statistical learning is impaired in individuals with developmental dyslexia, a specific learning disorder characterised by deficits in reading and spelling acquisition, despite normal intelligence and adequate school education (Lyon et al. 2003; Peterson and Pennington 2015). The worldwide prevalence of dyslexia is estimated at around 7 % (Yang et al. 2022), and the condition is often associated with drastic lifelong consequences for professional success, social inclusion, and emotional well-being, including higher rates of anxiety, depression, and suicide (Huntington and Bender 1993; Tsovili 2004; Carroll and Iles 2006). While originally regarded as a primarily phonological deficit (Vellutino et al. 2004), evidence has accumulated that dyslexia is associated with a broader and fundamentally perceptual deficit that affects individuals’ abilities to process (statistical) regularities in sensory input and to use these regularities to enhance the efficiency of auditory perception. The idea of such a broader deficit is supported by the fact that impaired perceptual processing was not only shown for speech stimuli, but also for non-linguistic sound materials. For instance, compared to healthy controls, individuals with dyslexia showed less behavioural benefits from stimulus repetition in perceptual tasks such as tone frequency discrimination (Ahissar et al. 2006; Gertsovski and Ahissar 2022) and less neural adaptation to different types of repeated auditory stimuli such as voices, tones (Perrachione et al. 2016; Jaffe-Dax et al. 2017; Jaffe-Dax et al. 2018; Peter et al. 2019), and phoneme categories (Ozernov-Palchik et al. 2022). In particular, processing of more complex statistical regularities was found to be impaired in dyslexia, including the extraction of transitional probabilities in sequences of syllables or tones (Gabay et al. 2015) and learning of distributional information such as frequencies of occurrence of certain syllables (Kimel et al. 2022) or bimodal distributions along a continuum between two non-native phonemes (Vandermosten et al. 2019). Using the statistical learning paradigm introduced above, one study showed reduced neural tracking at the rate of artificial words within a structured syllable stream in individuals with dyslexia, as well as a positive correlation between the strength of neural tracking at the word rate and phonological awareness (Zhang et al. 2021). Another recent study investigated the neural correlates of auditory statistical learning more generally in adults with and without dyslexia by passively presenting them with continuous sequences of (non-speech) sounds, i.e., tones with different frequencies, that were arranged into triplets (Daikoku et al. 2023). Occasionally, the last tone of a triplet was either a statistical deviant, i.e., it had a low transitional probability given the first two tones of the triplet, or an acoustic deviant, i.e., it was presented from a different location. In healthy adults, both types of rare deviant events elicited a mismatch response in the event-related potential (ERP), suggesting that they had formed expectations about upcoming tones based on the recent stimulus history, which were violated by the incoming deviant tone. By contrast, in adults with dyslexia, the mismatch response to acoustic deviants was diminished and no mismatch response was elicited by statistical deviants, indicating that listeners failed to extract transitional probabilities between tones to predict upcoming tones (Daikoku et al. 2023).

Various theoretical frameworks offer different explanations for the link between impaired statistical learning and phonological deficits in dyslexia (Singh and Conway 2021), including insufficient perceptual anchoring to previous stimuli (Ahissar 2007), impaired serial order learning (Szmalec et al. 2011), inefficient temporal sampling of continuous stimulation (Goswami 2011; Goswami 2019), or a reduced auditory memory span (Banai and Ahissar 2018). While the exact neurocognitive mechanisms remain poorly understood, it is plausible to assume that impaired statistical learning in dyslexia results in less robust phoneme representations as well as impedes the anticipation of upcoming events based on statistical regularities within a continuous speech stream, thereby reducing the efficiency of speech processing.

The aim of the present study was to gain a better understanding of how adults with dyslexia process statistical regularities in acoustic (non-speech) sequences, in comparison to healthy adults. While the recent EEG study introduced above (Daikoku et al. 2023) focussed on the automatic detection of rare deviant events that violate predictions based on transitional probabilities between tones within the continuous stream, the present study sought to investigate how auditory statistical regularities are neurally tracked during (passive) listening. To this end, we reanalysed the EEG dataset by Daikoku and colleagues (2023), using the approach by Batterink and Paller (2017) to quantify sensitivity to the statistical triplet structure in the tone sequence based on ITC of oscillatory neural activity. We hypothesised that the quotient of ITC at the rate of triplets relative to ITC at the rate of single tones, i.e., an index corresponding to the WLI in the study by Batterink and Paller (2017), would be reduced in adults with dyslexia compared to the healthy control group. This would indicate that sensitivity to the statistical structure of auditory sequences is lower, and regularities are less closely neurally tracked in dyslexia. In particular, weaker neural tracking of non-speech sound sequences (instead of syllable sequences as used by a previous study; Zhang et al. 2021) would provide support for the idea of a broader perceptual rather than a merely phonological deficit. Moreover, we predicted that neural tracking of the statistical triplet structure would be associated with individual spelling and reading scores as assessed using standardised tests prior to the experiment, such that individuals who show stronger neural tracking also achieve higher spelling and reading scores.

## Materials and Methods

### Participants

A total of 36 adult participants took part in the study. 17 of them (twelve female, five male; age M ± SD: 23.8 ± 4.2 years) had a diagnosis of developmental dyslexia, whereas the remaining 19 (13 female, six male; age M ± SD: 25.6 ± 3.3 years) formed the healthy control group. All participants had normal intelligence (IQ > 70; as assessed with a non-verbal intelligence test: Standard Progressive Matrices; Raven and Court 1998), no history of neurological or audiological disorders, and no diagnosis of general or specific language impairment. All of them were right-handed (as assessed with the Edinburgh Handedness Inventory; Oldfield 1971), native speakers of German, and had received a maximum of five years of formal musical training (besides music lessons at school). Approval of the experimental procedures was obtained from the local ethics committee (reference number: 2018/352), prior to the study. Participants provided written informed consent before the start of the study and received monetary compensation for their participation.

Before the actual EEG experiment, participants’ spelling and reading skills were assessed in a separate session to ensure that the groups indeed differed significantly in spelling and reading performance. As expected, members of the dyslexia group obtained lower scores than members of the control group in the German “Rechtschreibtest” (RST-ARR; Ibrahimović and Bulheller 2013), which required them to fill in missing irregular German words in a text that was read out by a native German person (*t*(34) = 9.71, *p* < .001, *d* = 3.373). Moreover, the dyslexia group showed reduced reading speed (*t*(34) = 5.29, *p* < .001, *d* = 1.766) and reading comprehension (*t*(34) = 4.91, *p* < .001, *d* = 1.637) compared to the control group, as assessed with the German “Lesegeschwindigkeits-und Verständnistest für die Klassen 5-12“ (LVGT 5-12+; Schneider et al. 2017). This test required them to read as much as possible of a continuous text within four minutes and to select from given words which of them fit some of the read text passages best.^1^

### Stimuli

The present study used the same sound stimuli as a previous study (Tsogli et al. 2019). A total of six Shepard tones based on different frequencies (F_3_ [174.61 Hz], G_3_ [196.00 Hz], A_3_ [220.00 Hz], B_3_ [246.94 Hz], C♯_4_ [277.18 Hz], and D♯_4_ [311.13 Hz]) were created by superpositioning nine sinusoidal components that were one octave apart. Each of the Shepard tones was combined with the sound of a percussion instrument (surdo, tambourine, agogô bells, hi-hat, castanets, and woodblock), with specific combinations of Shepard tones and percussion instruments counterbalanced across participants. For simplicity, these tones will be called tones A, B, C, D, E, F, and G in the remainder of the text. A separate set of six Shepard tones based on different frequencies was created for a short training phase prior to the start of the actual experiment. Additionally, one Shepard tone (based on C♯_5_ [554.37 Hz]) that was not combined with a percussive sound served as a target tone, which participants were asked to react to with a key press (see below). All tones had a duration of 220 ms, including a 10-ms rise and a 20-ms fall time, and a sampling frequency of 44,100 Hz.

The six tones (i.e., tones A, B, C, D, E, F) were arranged into two triplets (e.g., tones ABC and DEF; specific tone combinations were counterbalanced across participants), which were in turn combined to form a continuous tone sequence. Each 220-ms tone was followed by a 80-ms silent inter-stimulus interval, i.e., both within and between triplets, resulting in a triplet duration of 900 ms and a constant tone rate of 3.33 Hz for the whole sequence. Thus, the transitional probabilities between individual tones were the only cue available to segment the continuous tone sequence (at the triplet rate of 1.11 Hz). Two types of deviants occurred occasionally within the tone sequence: The last tone of a triplet was either replaced by the last tone of the respective other triplet in 10 % of the trials (statistical deviants), or was presented from a different speaker location in 20 % of the trials (acoustic deviants). Both types of deviants occurred also combined as double deviants in 2 % of all triplets. Since the aim of the present study was to investigate how listeners track statistical regularities in continuous auditory streams, deviants were not relevant to answer the research question of interest and only standard triplets without any deviant were included in the analysis.

### Procedure

Tone sequences were presented in six separate experimental blocks, each of which comprised 400 tone triplets and lasted approximately six minutes. During sound presentation, participants were seated inside an electromagnetically shielded chamber and EEG was recorded. Prior to the first experimental block, participants completed a short training phase (one minute) during which they could familiarise themselves with the task of pressing a key as fast as possible whenever they heard the target tone (i.e., the Shepard tone without a percussive sound). A total of 48 sounds were randomly replaced by the target tone throughout the experiment (eight per block), with target tone occurring equally often at each position within a triplet. To ensure that tracking of the (statistical) regularities of the tone sequence remained implicit, participants were not informed about the occurrence of deviants. In addition to the target detection task, they watched a silent documentary movie, which further diverted their attention away from the statistical regularities of the tone stream and reduced the probability that they focussed on the triplet structure between the rare target occurrences.

### EEG Data Acquisition

Continuous EEG was recorded from 64 scalp electrodes mounted in an elastic cap in accordance with the extended international 10-20 system. In addition to the scalp electrodes, horizontal and vertical electrooculograms were recorded bipolarly from electrodes placed at the outer canthus of each eye and above and below the right eye. The electrode on the left mastoid served as an online reference. During preparation, electrode impedances were kept below 5 kΩ. Signals were amplified with a BrainAmp amplifier (Brain Products, Munich, Germany), online band-pass filtered between 0.25 and 1000 Hz, and digitised with a sampling rate of 500 Hz.

### Data Analysis

EEG data were processed offline using the EEGLAB toolbox (version 2024.0; Delorme and Makeig 2004) in Matlab (version R2024a; The MathWorks, Natick, MA). Subsequent statistical analyses were done in RStudio (version 4.2.2, RStudio Inc., USA).

### Pre-processing

Before pre-processing, noisy channels whose signal variance exceeded an absolute z-score of 3.0 were excluded and later spherically spline interpolated. Continuous data of the remaining channels were first high-pass and then low-pass filtered with Kaiser-windowed finite impulse response filters at 0.2 Hz (transition bandwidth: 0.4 Hz, maximum passband deviation: 0.001, filter order: 4638) and 30 Hz (transition bandwidth: 5 Hz, maximum passband deviation: 0.001, filter order: 372), respectively. To remove physiological and technical artefacts, we used an independent component analysis (ICA). ICA was computed on a copy of the data that was filtered with a 1-Hz high-pass filter (transition bandwidth: 0.5 Hz, maximum passband deviation: 0.001, filter order: 3710) to improve signal-to-noise ratio for decomposition (with the same 30-Hz low-pass filter as above), and down-sampled to 125 Hz to shorten computation time. We obtained ICA weights with an infomax algorithm implemented in EEGLAB’s runica function and transferred them to the dataset that was pre-processed with the final filter parameters. Artefactual components were identified and selected for removal using a combined automatic and manual procedure: Independent components were first classified using the IC Label plugin for EEGLAB (Pion-Tonachini et al. 2019), and components that were classified as eye blinks or movements, muscle activity, cardiac activity, line noise, or channel noise with a probability of at least 30 % were automatically marked for rejection. Second, components classified as “other” were visually inspected with regard to their topography and spectral power distribution, and additionally selected for rejection if they were manually classified as one of the above-mentioned artefacts.

### Inter-Trial Coherence (ITC) and Triplet Learning Index (TLI)

Neural tracking of statistical regularities, i.e., the triplet structure, in the continuous tone sequences was assessed using inter-trial coherence (ITC) of brain responses.^2^ As already mentioned above, triplets that contained any type of deviant (statistical, acoustic, or double) were excluded from the analysis. Furthermore, triplets that immediately followed a deviant triplet as well as triplets that contained a behaviourally relevant target tone were discarded. Data of the remaining standard triplets were re-referenced to the algebraic mean of both mastoid electrodes. To attenuate baseline differences within and between triplets, we first extracted epochs that ranged from 0 to 300 ms relative to tone onset, i.e., covered one tone plus the silent inter-tone interval, and de-meaned them individually. For the computation of ITC, we then generated longer non-overlapping epochs with a duration of 5.4 sec, each of which covered six triplets (i.e., 18 de-meaned epochs of individual tones). Epoching was done separately within each experimental block to make sure that no epochs were generated that contained block boundaries (corresponding to breaks within the experimental session). Any 5.4-sec epoch with a peak-to-peak difference that exceeded 300 µV was discarded from the analysis, resulting in an average of 32 ± 1 (M ± SD) epochs per block for each participant. To improve signal-to-noise ratio, these data were averaged across nine electrodes within a fronto-central cluster (F1, F2, Fz, FC1, FC2, FCz, C1, C2, Cz), in line with the topographies shown by similar previous studies (e.g., Batterink and Paller 2017). Separately for each block, epochs were multiplied with a Hanning window to reduce 1/f noise, before Fast Fourier transforms (FFTs) were computed. In accordance with recent recommendations on how to best quantify neural tracking of sensory input (Benjamin et al. 2021), results of the FFT were averaged before ITC was estimated from the extracted phase angles. ITC coefficients were then extracted at the tone (3.33 Hz) and triplet (1.11 Hz) rate to compute the Triplet Learning Index (TLI) as the quotient of the ITC coefficient at the triplet rate divided by the ITC coefficient at the tone rate, corresponding to the Word Learning Index reported by Batterink and Paller (2017). The TLI is thought to reflect sensitivity to the triplet structure of the sequence, assuming that a shift towards relatively higher ITC at the triplet rate and relatively lower ITC at the tone rate over time indicates learning of the statistical regularity. We used a one-sided independent-samples *t*-test to statistically test the hypothesis that the dyslexia group shows a lower TLI (averaged across experimental blocks) compared to the control group, i.e., a weaker sensitivity to the statistical triplet structure.

To illustrate how the TLI changes as a function of statistical learning of the triplet structure over the course of the first block (during which most learning is expected to occur), we used a sliding time window approach (for a similar approach, see Batterink and Choi 2021). Therefore, we computed ITC in windows of 30 consecutive non-overlapping 5.4-sec epochs (i.e., window length = 162 sec), and shifted these windows in steps of 5.4 sec. ITC was plotted as a function of the number of triplets presented so far. We then fitted a logarithmic function (y = a * log(x) + b) to the ITC time course, separately for the dyslexia and the control group.

Moreover, we re-computed ITC for every single electrode (instead of the fronto-central cluster) to visualise the topographical distribution of ITC (on average across experimental blocks) at the triplet and syllable rate, respectively.

### Brain-Behaviour Correlations

Finally, we computed brain-behaviour correlations to assess the relationship between neural tracking of the statistical triplet structure in the tone sequences and spelling and reading performance (as assessed in the standardised tests explained above). To this end, raw test scores from the spelling test (RST-ARR) as well as raw scores of reading speed and reading comprehension from the reading test (LVGT 5-12+) were z-standardised. We then computed three separate Pearson correlations between each of the scores and the mean TLI across experimental blocks, respectively, and one-sided single-sample *t*-tests were used to test whether the correlation coefficients were significantly above zero.

## Results

As displayed in Figure 1A, the largest peak in ITC in the fronto-central electrode cluster (across all experimental blocks) was visible at the tone rate (3.33 Hz) in both the control and the dyslexia group. Clear peaks in ITC also occurred at the triplet rate (1.11 Hz), as well as at its harmonics (2.22 Hz and 4.44 Hz). ITC at the tone rate was lower for the control group compared to the dyslexia group, whereas ITC tended to be slightly higher for the control group compared to the dyslexia group at the triplet rate (and its harmonics). The ITC effect at both tone rate and triplet rate was broadly fronto-centrally distributed across both groups, as shown in Figure 1B, consistent with an origin in temporal (auditory) cortex. Thus, in both groups, participants’ brain activity became synchronised with the tone rate in the stimulation, and all participants picked up the statistical triplet structure in the tone sequences to some extent.

**Figure 1.**
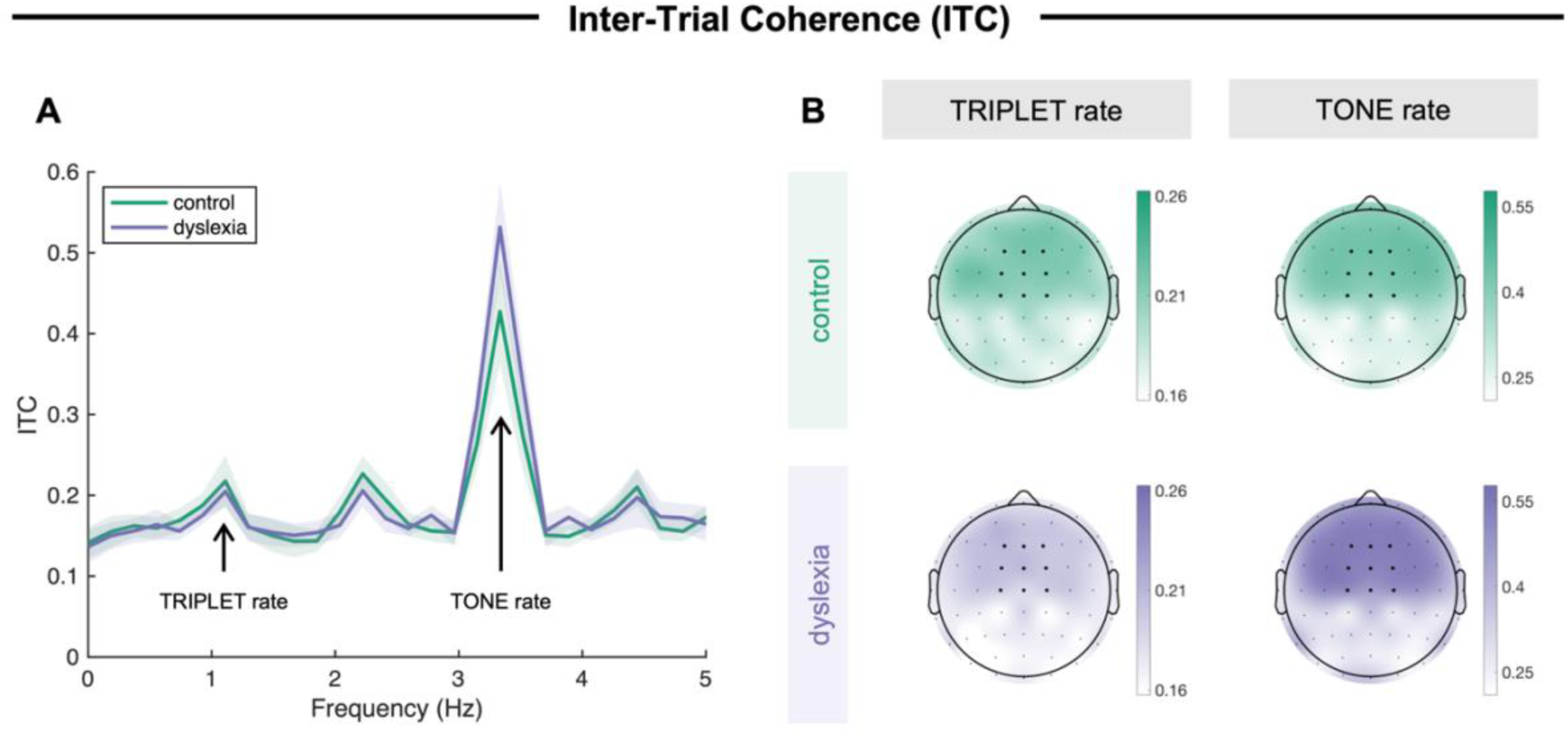
A: Inter-trial coherence of neural activity (y-axis) across frequencies (x-axis) in the fronto-central electrode cluster of interest for the control and the dyslexia group. As marked in the plot, 3.33 Hz corresponds to the tone rate, and 1.11 Hz corresponds to the triplet rate. Shaded areas indicate ± 1 standard error of means. B: Topographies of ITC at the triplet rate (left panels) and the tone rate (right panels) for the control (upper panels) and the dyslexia (lower panels) group. Please note that different scales were used for the triplet and the tone rate. Black dots in the topography plots indicate electrode locations, and bold dots mark electrodes included in the fronto-central electrode cluster of interest.

Crucially, the TLI (displayed in Figure 2) revealed that the relative weighting of synchronised activity at the tone rate and the triplet rate differed between the control and the dyslexia group: On average across experimental blocks, the TLI was significantly higher in the control group than in the dyslexia group (*t*(33) = 1.88, *p* = .034, *d* = 0.64)^3^, suggesting a relatively higher sensitivity to the triplet structure of the sequence (see Figure 2A). As shown in Figure 2B, this difference between groups emerged early throughout the first block, when TLI increased more steeply in control participants than in individuals with dyslexia as the number of presented triplets increased.

**Figure 2.**
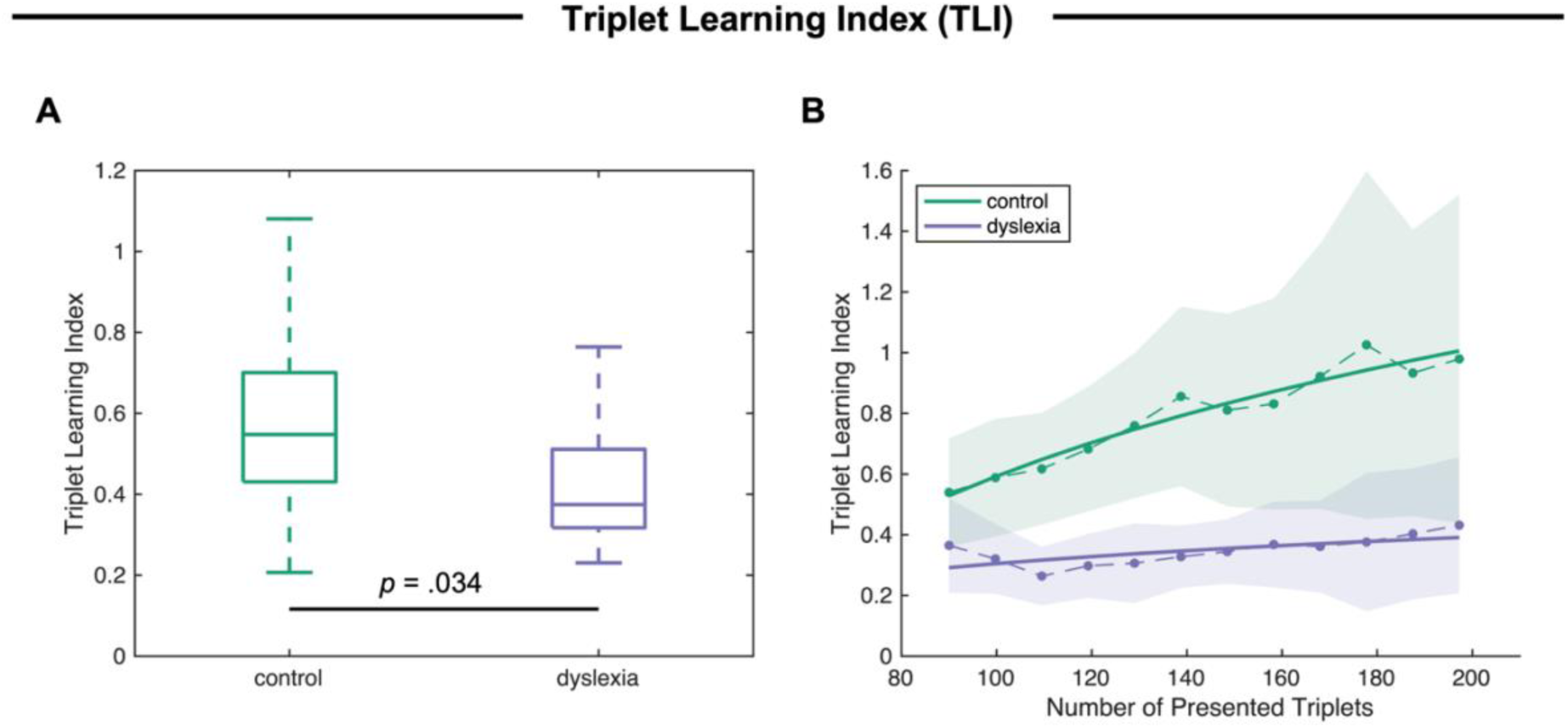
A: Mean Triplet Learning Index across experimental blocks for the control and the dyslexia group. Boxes indicate the inter-quartile range (around the median line), and whiskers indicate minimum and maximum per group. B: Triplet Learning Index (y-axis) as a function of the number of presented triplets (x-axis) throughout the first experimental block for the control and the dyslexia group. Dashed lines correspond to the actual data, and bold solid lines are logarithmic functions (y = a * log(x) + b) fitted to the data. Shaded areas indicate ± 1 standard error of means (in the actual data).

**Figure 3.**
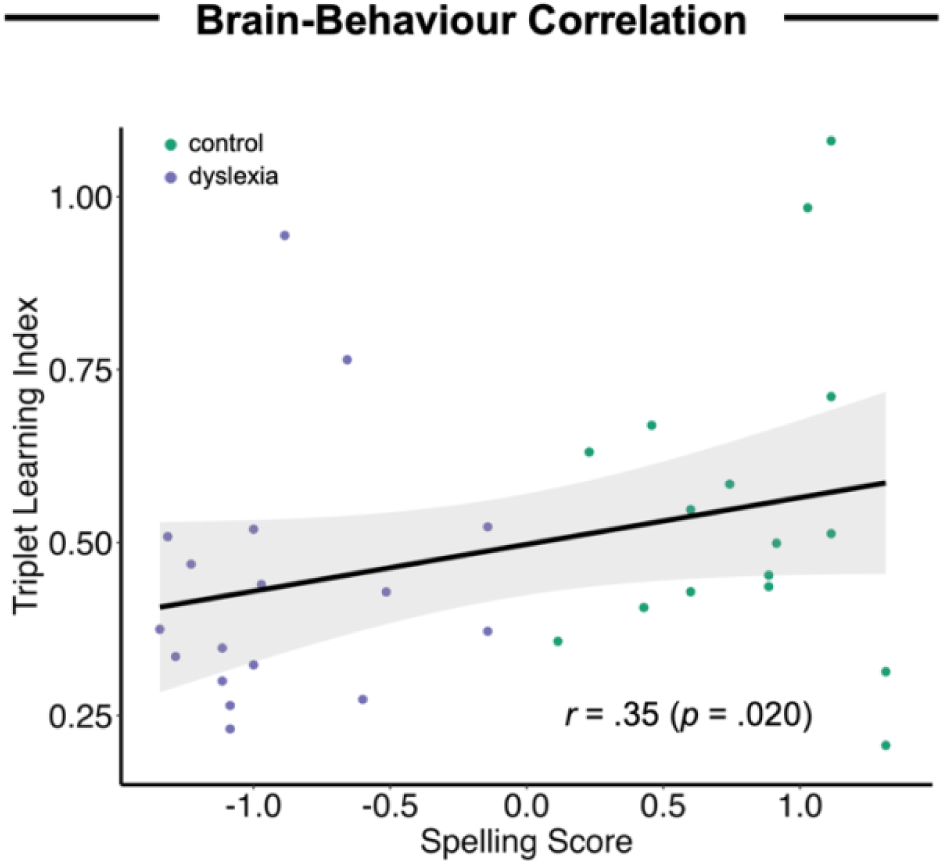
Correlation between the individual spelling scores (x-axis) and Triplet Learning Index (y-axis). Shaded areas around the trend line indicate the 95-% confidence interval.

Finally, we found a significant positive correlation between participants’ TLI across experimental blocks and their standardised score in the spelling test (*r* = .35, *p* = .020). This suggests that participants with a higher spelling performance also showed stronger neural tracking of the statistical triplet structure in the tone sequence. No significant relationship was found between participants’ TLI and their reading speed (*r* = .01, *p* = .478) and reading comprehension (*r* = -.06, *p* = .624). The absence of a significant correlation for reading scores suggests that the positive correlation between participants’ TLI and spelling scores is not merely driven by group selection.

## Discussion

The current study aimed to compare neural tracking of the statistical structure of a sequence of tones between adults with and without dyslexia. Two groups of participants were passively presented with a continuous stream of tones, which were arranged into triplets, such that the transitional probabilities between tones were high within a triplet and low between triplets. Sensitivity to the statistical triplet structure was quantified with the TLI, i.e., the ratio of ITC at the triplet rate divided by ITC at the tone rate. While both groups did show an ITC peak at the triplet rate, suggesting that they picked up the triplet structure to some extent, the TLI was substantially lower in the dyslexia than in the control group. This indicates that individuals with dyslexia were relatively less sensitive to the statistical structure of the tone sequence, and their brains tracked the triplet rhythm less closely relative to the syllable rate. The difference between groups emerged early throughout the first experimental block, when the TLI increased much more steeply for healthy adults than for adults with dyslexia, pointing towards rapid learning of statistical regularities in the control, but not in the dyslexia group. Furthermore, the TLI was significantly correlated with individual spelling scores, such that stronger neural tracking of the triplet structure was associated with better spelling skills, but not with reading scores. The significant correlation suggests that the ability to acquire statistical regularities from the auditory environment is linked with (at least some) literacy skills that are relevant for the clinical diagnosis of dyslexia.

Overall, these results are in line with a large body of previous research that demonstrated impaired processing of (statistical) regularities in individuals with dyslexia across age groups. Earlier studies reported smaller behavioural benefits of stimulus repetition (Ahissar et al., 2006), diminished neural adaptation to different types of linguistic and non-linguistic sound materials (Perrachione et al. 2016; Jaffe-Dax et al. 2017; Jaffe-Dax et al. 2018; Peter et al. 2019; Ozernov-Palchik et al. 2022), reduced learning of more complex statistical regularities, including transitional probabilities (Gabay et al. 2015; Daikoku et al. 2023) and distributional statistics (Vandermosten et al. 2019; Kimel et al. 2022), and diminished neural tracking of the word structure within a syllable stream (Zhang et al. 2021). The current study extends these findings by comparing online neural tracking of statistical regularities in sequences of non-speech sounds between individuals with and without dyslexia. Specifically, the present results are in line with the hypothesis that the failure to detect rare, unexpected (statistical) deviant events that was observed in adults with dyslexia (Daikoku et al. 2023) may stem from their difficulties in tracking the regular statistical structure of the sensory input.

Our finding of weaker relative tracking of statistical regularities in individuals with dyslexia has several implications both for theories on the neuro-cognitive underpinnings of developmental dyslexia and for possible clinical applications. From a theoretical point of view, the present results argue against a specific phonological deficit as the main cause underlying dyslexia (Vellutino et al. 2004). Instead, our findings support the idea of a somewhat broader perceptual impairment in processing statistical structure in sound. In particular, the presence of an effect despite the use of non-linguistic stimulus material indicates that the statistical learning deficit in individuals with dyslexia is not limited to speech input, but affects auditory processing more generally across a wider range of acoustic stimuli. Moreover, the correlation between neural tracking of the statistical structure of the tone sequence and individual spelling scores suggests that these abilities are interconnected. This relationship may be explained via phonological representations, which seem to be atypical in individuals with dyslexia: There is evidence that the perception of phonemes is less consistently categorical in both adults (Vandermosten et al. 2010) and children with dyslexia (Vandermosten et al. 2011) than in healthy control participants when they are asked to behaviourally classify stimuli along a synthetic phoneme continuum. Impoverished representations of phoneme categories may be grounded in deficits in statistical learning, which has been argued to play a crucial role for the acquisition of novel phoneme categories. More specifically, listeners are thought to acquire knowledge about the distribution of acoustic features through exposure to natural language, e.g., they learn that certain features (e.g., voice onset time, which distinguishes voiced and voiceless stop consonants such as /d/ and /t/) are not randomly distributed across a continuum, but cluster around certain values to characterise and distinguish different phonemes (Myers 2014). Deficits in learning such distributional statistics, as they were reported for individuals with dyslexia, may result in impoverished representations of phoneme categories (Banai and Ahissar 2018), which in turn weaken phonological awareness, i.e., an individual’s knowledge about the sound structure of speech, which is an essential skill for reading and spelling development (Snowling 2000; Ziegler and Goswami 2005).

We can only speculate about why statistical learning was only correlated with individual spelling scores, but not with reading scores in the present dataset; however, it may be plausible to assume that adults have developed compensatory strategies such as the use of semantic context information, which is typically available during reading, but does not help during spelling. In the previous analysis of ERPs to statistical deviants in the same dataset, no correlation was found between ERP amplitudes and reading or spelling skills (Daikoku et al. 2023). The presence of a significant correlation of neural tracking with spelling skills in the present study suggests that continuous neural tracking is a more sensitive and direct measure of statistical structure learning than ERP responses to rare, unexpected statistical deviants.

From a more clinical and applied point of view, our present results suggest that statistical learning paradigms may be useful as an additional diagnostic tool in individuals with dyslexia or a screening tool in individuals at risk of dyslexia. The statistical learning paradigm has yielded robust results in children (Saffran et al. 1997), infants (Saffran et al. 1996), and even neonates (Teinonen et al. 2009), and here we show that neural correlates of statistical learning of non-linguistic auditory stimuli are related to participants’ spelling skills. Thus, such paradigms may be well-suited as a screening tool because they are relatively simple and efficient, do not require active behavioural responses, and do not rely on linguistic stimulus material; therefore, they may also be used in pre-verbal children, people with language impairments, and across different languages.

In conclusion, the present study demonstrated that neural tracking of statistical regularities in auditory (non-speech) stimulus sequences is reduced in adults with dyslexia compared to healthy adults, and that the magnitude of neural tracking of the statistical stimulus structure is associated with individual spelling scores. By combining an online measure of neural tracking of stimulus regularities and a statistical learning paradigm with non-linguistic sound material in individuals with dyslexia, these results both provide empirical support for theories claiming a link between statistical learning abilities and literacy skills and point out the potential use of such paradigms as a screening tool for dyslexia. To further explore the link between perceptual processing of statistical structure in sound and phonological skills and to derive new intervention approaches to ameliorate phonological deficits in dyslexia, future research may include groups of participants who are expected to show particularly good statistical learning abilities through extensive training, such as experienced musicians (Pesnot Lerousseau and Schön 2021), and involve longitudinal designs with different age groups across development.

## Acknowledgements

This research was supported by a Moonshot Goal 9 grant by the Japan Science and Technology Agency (JST) awarded to T.D. (grant number JPMJMS2297) and a Postdoctoral Fellowship for Research in Japan by the Japan Society for the Promotion of Science (JSPS) awarded to H.R. (grant number PE24035). The funding sources had no role in the design of this study, data analysis and interpretation, and the decision to publish this manuscript.

1 For further details on the spelling and reading scores, please refer to the original publication of the dataset (Daikoku et al. 2023).

2 As a complementary measure, we also computed spectral magnitude. Spectral magnitude showed a virtually identical pattern of peaks across frequencies as did ITC, as expected. Therefore, we chose to report the ITC results in detail, since such as phase-based measure is less likely to depend on amplitude changes of the signal than power-based measures (Benjamin et al. 2021).

3 One participant from the control group was excluded from the analysis because their TLI differed from the group mean by more than 4 SD. Thus, the reported statistical results refer to data from 18 control participants and 17 individuals with dyslexia.

